# Molecular determinants of vascular transport of dexamethasone in COVID-19 therapy

**DOI:** 10.1101/2020.07.21.212704

**Authors:** Ivan G. Shabalin, Mateusz P. Czub, Karolina A. Majorek, Dariusz Brzezinski, Marek Grabowski, David R. Cooper, Mateusz Panasiuk, Maksymilian Chruszcz, Wladek Minor

## Abstract

Dexamethasone, a widely used corticosteroid, has recently been reported as the first drug to increase the survival chances of patients with severe COVID-19. Therapeutic agents, including dexamethasone, are mostly transported through the body by binding to serum albumin. Herein, we report the first structure of serum albumin in complex with dexamethasone. We show that it binds to Drug Site 7, which is also the binding site for commonly used nonsteroidal anti-inflammatory drugs and testosterone, suggesting potentially problematic binding competition. This study bridges structural findings with our analysis of publicly available clinical data from Wuhan and suggests that an adjustment of dexamethasone regimen should be considered for patients affected by two major COVID-19 risk-factors: low albumin levels and diabetes.

**One Sentence Summary:** Structure of serum albumin with dexamethasone reveals why the drug may not always help COVID-19 patients.

## Introduction

On June 16 2020, collaborators in the RECOVERY Trial (Randomized Evaluation of COVID-19 Therapy) reported that a daily dose of 6 mg dexamethasone reduced deaths among COVID-19 patients receiving respiratory interventions (*1, 2*). This widely available steroid has been shown to cut deaths by approximately 30% for patients who were on ventilators and 20% for those who were receiving oxygen therapy but were not on ventilators. Dexamethasone is a corticosteroid exhibiting potent anti-inflammatory and immunosuppressant effects. Since the 1960s, it has been used to treat many conditions, including severe pneumonia, rheumatic problems, skin diseases, severe allergies, asthma, chronic obstructive lung disease, brain swelling, and many others (*3, 4*). Dexamethasone suppresses the immune system by stimulating the glucocorticoid receptor, thus providing some relief for patients whose lungs are devastated by an overactive immune response that accompanies severe cases of COVID-19, but no benefit for those patients who did not require respiratory support (*5*). Along with anti-inflammatory action, steroids can also reduce airway spasm and obstruction in patients who have asthma or chronic obstructive pulmonary disease – groups of patients of especially high risk (*6*). In the RECOVERY trial, the therapeutic effect of dexamethasone appeared to depend on “using the right dose, at the right time, in the right patient” (*5*). However, there is another important factor which might have influenced the effectiveness of dexamethasone: its vascular transport.

Dexamethasone is primarily delivered throughout the body by serum albumin; about 77% of dexamethasone in blood is bound to plasma proteins, mostly to SA (*4*). Serum albumin has a typical blood concentration of 35-55 g/L (600 μM) and constitutes up to 55% of total plasma proteins (*7*). During its month-long lifetime, an albumin molecule makes nearly 15,000 trips around the body, facilitating vascular transport of hormones, metals, fatty acids, and drugs (*8*). The extensive drug-binding capacity of albumin is enabled by its high concentration and by the presence of at least 10 distinct drug-binding sites on the molecule (*9*). Preferred binding sites on albumin are known only for 32 drugs, including ibuprofen, warfarin, cetirizine, naproxen, diazepam, etodolac, and halothane (*9*). The binding capacity of albumin can be affected by several factors, and the inconsistent therapeutic action of dexamethasone on COVID-19 may be partially explained by differences in its vascular transport. Herein, we report the molecular structure of albumin in complex with dexamethasone, provide new insights into the mechanism of its transport, and discuss how compromised vascular transport of dexamethasone may limit its effectiveness in COVID-19 therapy.

### Structure of serum albumin in complex with dexamethasone

The crystal structure of equine serum albumin (ESA) in complex with dexamethasone was determined at 2.4 Å resolution (**Table S1**). The overall fold of albumin in this structure is essentially identical to previously published structures of human serum albumin (HSA) and ESA (**Table S2**). The electron density maps clearly indicate one dexamethasone molecule bound in Drug Site 7 (DS7), which is located in domain II, between subdomains IIA and IIB (**Fig. 1**) The drug site number is according to the recently expanded nomenclature (*9*), which was built upon previously used site names (*10, 11*).

**Figure 1.**
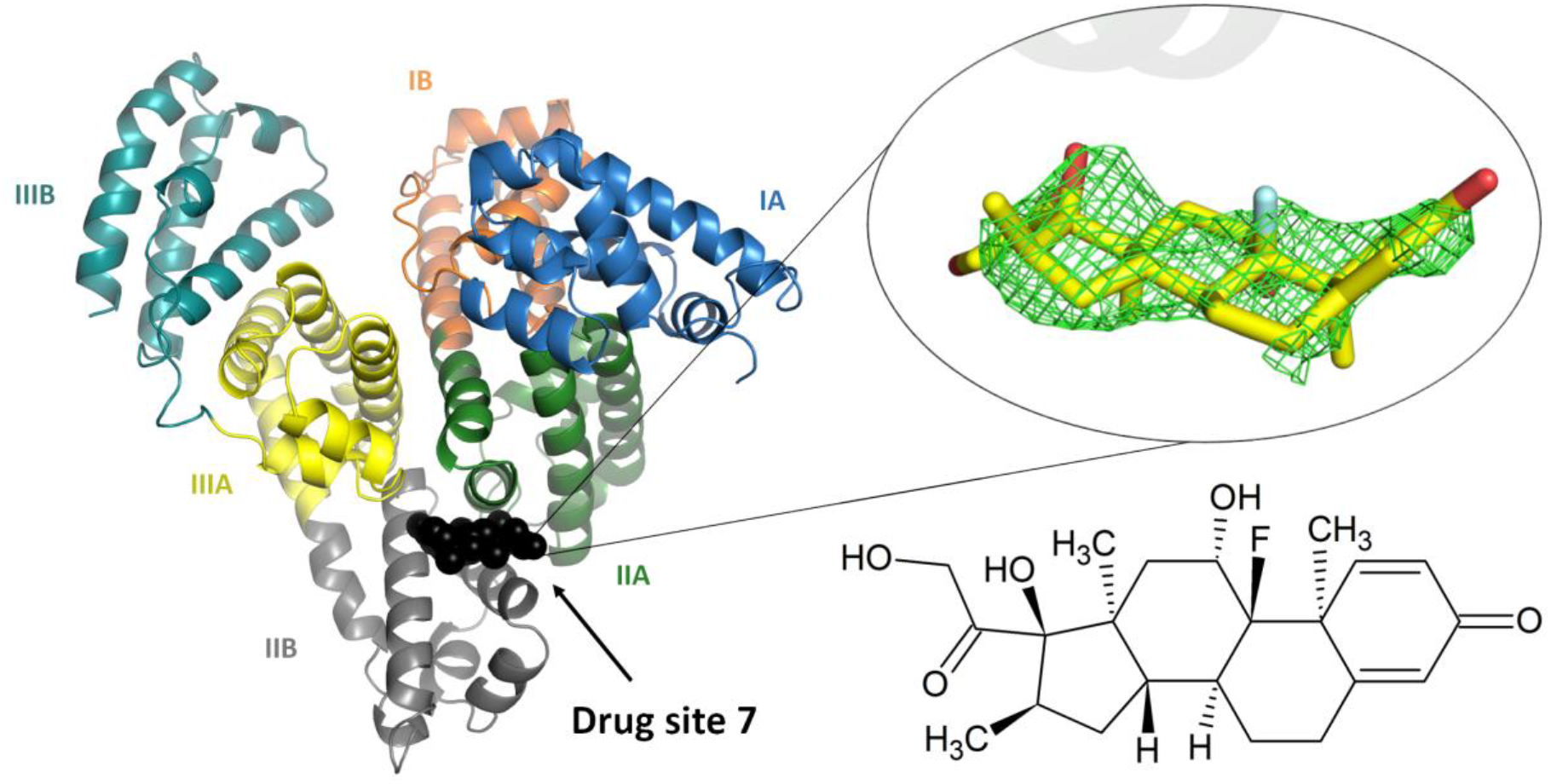
The overall structure of the ESA-dexamethasone complex. The electron density observed for dexamethasone in DS7 is shown as green mesh (mF_o_–DF_c_ map, calculated after ten refinement cycles without the ligand, RMSD 2.5). Albumin subdomains are shown in different colors and labeled with Roman numerals and letters (e.g., IA). The dexamethasone molecule is shown in stick representation with carbon atoms in yellow, oxygen atoms in red and fluoride atom in cyan. The chemical structure of dexamethasone is displayed in the same orientation as the stick representation. The electron density, including the omit maps, and the model can be inspected interactively at https://molstack.bioreproducibility.org/project/view/gmvG1L8c66YgPpgqS0Zm/

Fifteen residues are found within 5 Å of dexamethasone molecule (**Fig. S1**): Arg208, Ala209, Lys211, Ala212, Val215, Asp323, Leu326, Gly327, Leu330, Leu346, Arg347, Ala349, Lys350, Glu353, and Ala481 (corresponds to Val482 in HSA). Side-chains of residues Ala212, Val215, Leu326, Leu330, Leu346, Ala349, and the hydrophobic part of Lys350 form a mostly hydrophobic surface at the inner side of the binding cavity, towards which the most hydrophobic part of dexamethasone molecule is oriented (**Fig. 2A**). The cavity is partially separated from the solvent by a strong salt bridge between Arg208 and Asp323, which bridges the IIA and IIB subdomains. In addition to the hydrophobic interactions, the drug molecule is stabilized by two hydrogen bonds: O2 hydroxyl group with NH2 atom of Arg208 and O3 hydroxyl group with the main chain oxygen of Arg208. In comparison, dexamethasone forms six hydrogen bonds with the glucocorticoid receptor (*12*). The significantly higher number of hydrogen bonds to the target of its action (the glucocorticoid receptor) than to the transport protein (albumin) is reflected in the respective binding constants: dexamethasone dissociation constant to HSA is 58.8 μM (*13*), which is four orders of magnitude higher than that to the glucocorticoid receptor – 4.6 nM (*14*).

**Figure 2.**
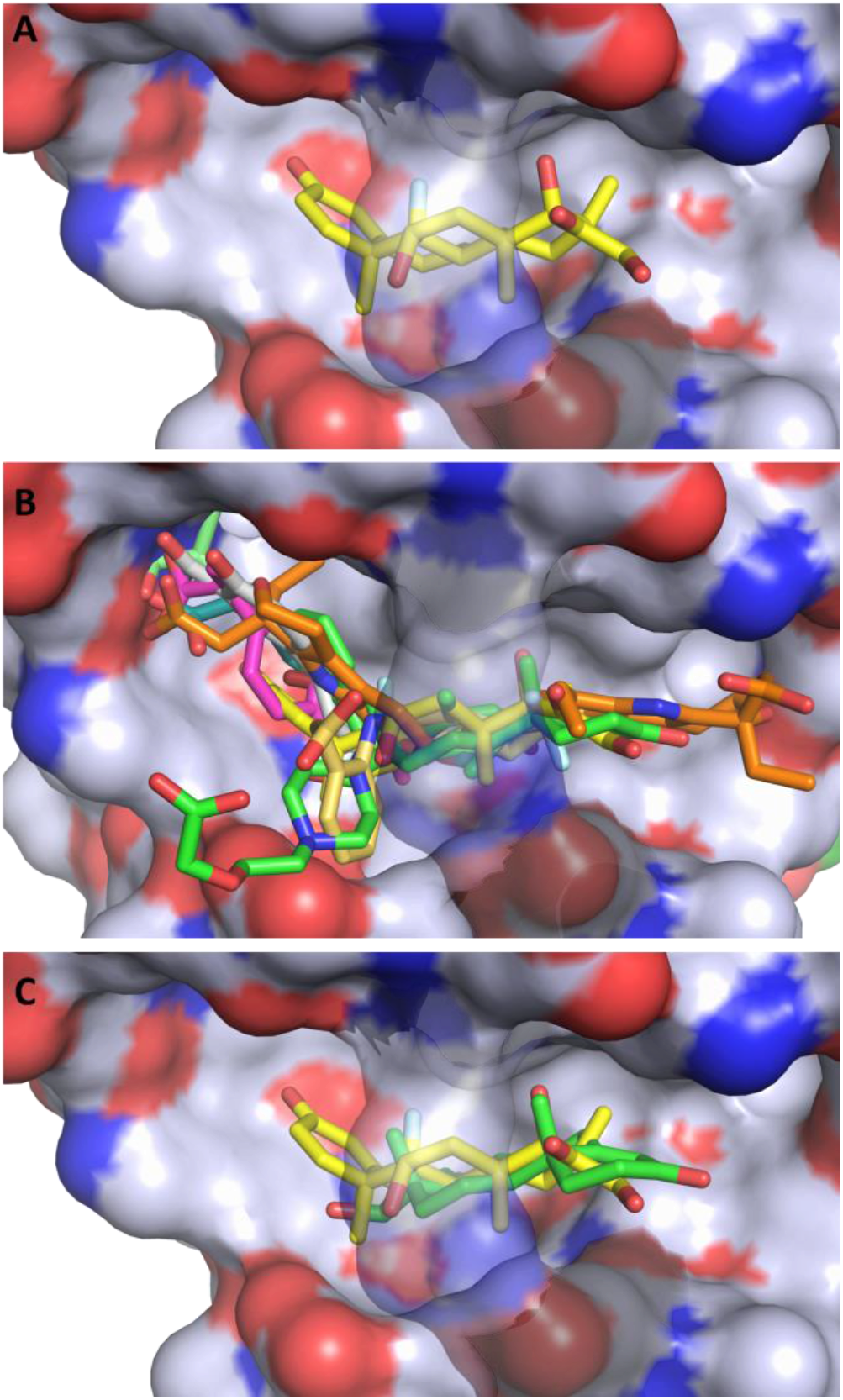
Hydrophobicity of Drug Site 7 and potential competition between dexamethasone and other drugs. (**A**) Dexamethasone bound to ESA. The color scheme for the protein surface is as follows: gray for the contribution from carbon atoms, red for oxygen atoms, and blue for nitrogen atoms. The link covering the cavity is formed by salt bridge between Arg208 and Asp323 and is transparent on all panels for clarity. (**B**) Superposition of the serum albumin complexes with all of the FDA-approved drugs known to bind to DS7: dexamethasone (PDB ID: 6XK0), ibuprofen (PDB ID: 2BXG), diflunisal (PDB ID: 2BXE), cetirizine (PDB ID: 5DQF), testosterone (PDB ID: 6MDQ), halothane (PDB ID: 1E7B), naproxen (PDB ID: 4ZBR), 6-MNA (PDB ID: 6U5A), diclofenac (PDB ID: 6HN0), and etodolac (PDB ID: 5V0V). (**C**) Superposition of the ESA-dexamethasone (PDB ID: 6XK0) and ESA-testosterone (PDB ID: 6MDQ) structures. Dexamethasone and testosterone largely overlap at DS7. Both steroids are shown in stick representation with oxygen atoms in red; dexamethasone is shown with carbon atoms in yellow and fluoride atom in cyan, while testosterone is shown with carbon atoms in green.

A high sequence identity/similarity between HSA and ESA (76.1%/86.2%) results in similar drug-binding properties of these albumins. Moreover, DS7 is classified as one of the most conserved drug-binding sites between ESA and HSA (*9*). For example, the residue conservation in this site was shown to result in similar testosterone-binding properties of this site in albumin from both species (*15*). Similarly, 14 out of 15 residues involved in dexamethasone binding to ESA are conserved in HSA (**Fig. S1**). Only one residue is different: Ala481 in ESA corresponds to Val in HSA. This small hydrophobic-for-hydrophobic difference is unlikely to affect dexamethasone binding in HSA because it is a kind-for-kind change that does not introduce any clashes with the ligand (**Supplementary text**). Therefore, the conservation of amino acid residues in DS7, which leads to essentially identical hydrophobic environments, suggests that dexamethasone binds to HSA in the same site as in ESA. This conclusion is supported by the previously reported spectroscopic studies of dexamethasone binding by HSA and BSA, which predicted that dexamethasone binds to the hydrophobic pocket around Trp214 inside the IIA subdomain (*13*). In our ESA-dexamethasone structure, the distance between dexamethasone and the indole ring of Trp214 is about 12 Å.

### Competing compounds and albumin glycation may impair dexamethasone binding

Drug Site 7 has been structurally characterized for human, equine, bovine, ovine, caprine, and leporine albumin, and has been shown to bind testosterone, several nonsteroidal anti-inflammatory drugs (diclofenac, diflunisal, etodolac, ibuprofen, naproxen, and 6-MNA), cetirizine, and the general anesthetic halothane (*9*). The site is formed by a large cavity, and some of the drugs shown to bind there occupy slightly different, yet overlapping, locations within the site (**Fig. 2B**). Multiple drugs binding in the same binding site may result in drug-drug displacement: a drug may be displaced by another drug administered at a high concentration. The increase in the free fraction of the displaced drug results in its decreased half-life and potential toxicity (*16, 17*), which is especially concerning for drugs whose margins of safety are small (*17*). Although albumin is a major drug transporting protein, structures of its complexes with only 32 FDA-approved drugs have been determined so far (*9*), and it is unknown which drugs used in COVID-19 treatment would compete with dexamethasone. For example, colchicine (Colcrys), one of the drugs tested for effectiveness in COVID-19 patients, and theophylline (Elixophyllin), which is used in asthma and chronic obstructive pulmonary disease, have narrow therapeutic ranges and in the blood bind to albumin in 40% (*18–20*) but their binding sites are unknown. Therefore, there is an urgent need for mapping albumin binding sites for most commonly prescribed drugs. Currently available structural data suggest potential competition between dexamethasone and drugs that bind to DS7 (**Fig. 2B**), some of which were already reported to affect the way dexamethasone works (*21, 22*).

Similarly, dexamethasone overlaps with the testosterone molecule (**Fig. 2C**), which is the only steroid for which a structure of a complex with albumin has been determined (*15*). Testosterone is suspected of playing a critical role in driving the pronounced excess of COVID-19 lethality in male patients (*23*), and low testosterone levels predict adverse clinical outcomes (*24*). Critically ill male COVID-19 patients suffer from severe testosterone deficiency, which may or may not be caused by the disease. Dexamethasone competition with testosterone for the same binding site may further exacerbate the effect of low testosterone by affecting its transport, especially if high doses of dexamethasone are administered.

Moreover, the location of dexamethasone binding site suggests that dexamethasone transport may be affected by glycation of albumin observed in diabetic patients. Diabetes is one of the major risk-factors in COVID-19: 25.7% of males and 20% of females among admitted patients in one study had type-II diabetes (*25*). From the perspective of blood drug transport, diabetes presents an additional challenge due to glycation of drug-binding sites on albumin. Glycation, which is significantly increased in diabetic patients due to the elevated blood glucose levels, alters albumin drug-binding ability. Four residues in DS7 can be glycated; among them, glycation of Arg208 is likely to prevent drug binding in this site due to potential steric clashes and disruption of the binding site by elimination of the Arg208-Asp323 salt bridge (**Supplementary text**), thus decreasing albumin binding capacity for dexamethasone.

### Albumin and glucose levels of COVID-19 patients

We have analyzed the albumin levels of COVID-19 patients admitted to Tongji Hospital, Wuhan, China between January 10 and February 18, 2020 (*26*). Out of 373 patients (222 males, 151 females), 174 died and 199 survived. The high mortality rate seen in this cohort (46.6%) stems from the fact that Tongji Hospital admitted a high rate of severe cases in Wuhan (*26*). The albumin levels for both outcome groups were first positively tested for normality using Q-Q plots (**Fig. S2**) and the Kolmogorov-Smirnov test (*W_Died_*=0.995, W_Survived_=0.991). The difference between mean albumin levels was found to be statistically significant according to Welch’s t-test (*p* < 0.001) and showed that patients that died were associated with much lower albumin levels (*μ_Died_*=27.8 g/L) than those that survived (*μ_Survived_*=36.8 g/L). The unadjusted odds ratio of surviving COVID-19 for albumin was 1.56 (95% CI: 1.44–1.71, *p* < 0.001), whereas the odds ratio adjusted for gender, age, and glucose level was 1.51 (95% CI: 1.37–1.69, *p* < 0.001); detailed odds ratios for both models are presented in **Table S3**. This means that each 1 g/L increase in the patient’s albumin level was associated with 50% increased odds of surviving COVID-19. Other studies (**Table S4**) linked low levels of albumin with poor outcomes in COVID-19 (*27–29*), but they did not provide supporting blood sample data that would allow full, time-series analyses of relations between different risk-factors and patient survival rate. **Figure 3** shows patient albumin levels in relation to the outcome, gender, time since admission, age, and glucose level.

**Figure 3.**
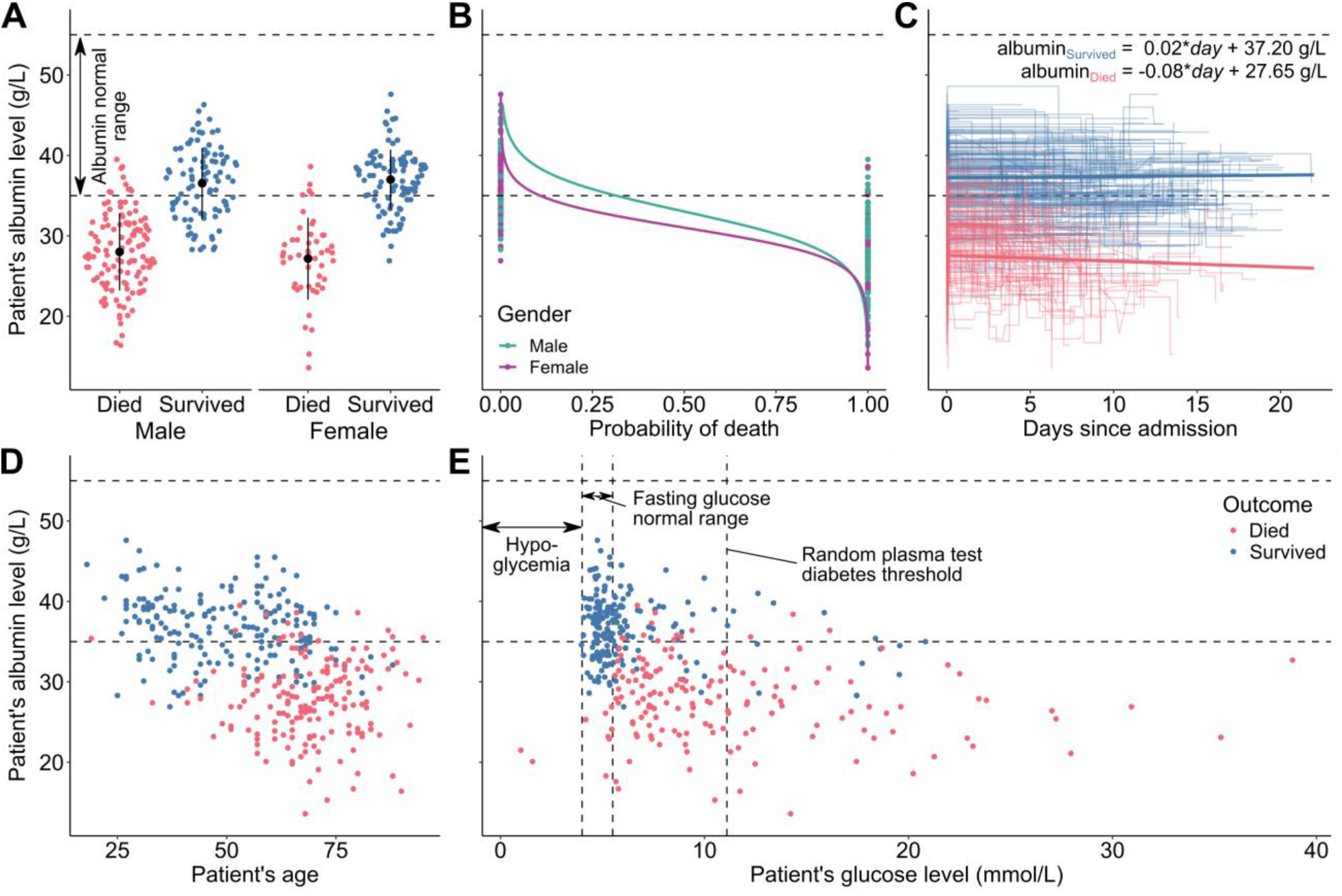
Albumin levels of 373 patients from Tongji Hospital, Wuhan, China, admitted between January 10 and February 18, 2020. (**A**) Violin strip charts of final albumin levels for patients that died (red) and survived (blue), grouped by gender; means and standard deviations overlaid. (**B**) Logistic regression showing the difference between the probability of death for men (teal) and women (purple), at different albumin levels. (**C**) Line plots presenting each patient’s albumin level over several blood samples taken during their time in the hospital; median linear regression line between the first and last blood samples for each outcome group overlaid. (**D**) Scatter plot presenting the relation between albumin level and age. (**E**) Scatter plot presenting the relation between final albumin and glucose levels. Source data from (*26*).

The majority of patients that died of COVID-19 had albumin levels not only lower than the patients that survived but also below the normal range, which is usually defined as 35-55 g/L (*7*). There is also a difference between the mortality rate among males (56.8%) and females (31.8%). Even though albumin level distributions for both genders have similar shapes, the proportions of survivors were not the same (**Fig. 3A**). The adjusted odds ratio for surviving if a patient was female was 2.19 (95% CI: 1.04–4.72, *p* < 0.05), however the differences in survival chances corresponded mainly to high albumin levels (**Fig. 3B**). This means that very low albumin levels were a strong predictor of death, regardless of gender. Moreover, the albumin levels of patients varied during their hospital stay (**Fig. 3C**). The correlation between admission and outcome levels was statistically significant only for patients that died (*r_Died_*=-0.22, *p* < 0.001; *r_Survived_*=-0.07, *p* = 0.176), which indicates that patients that died from COVID-19 suffered a decrease of albumin levels during their hospital stay. Furthermore, age was also a risk factor (**Fig. 3D**), with each year decreasing the odds of survival by 8% (odds ratio: 0.92, CI 95%: 0.89–0.94, *p* < 0.001). Finally, it can be noticed that high glucose levels were also a significant risk factor (**Fig. 3E**). The adjusted odds ratio of surviving COVID-19 for glucose was 0.89 (CI 95%: 0.81–0.96, *p* < 0.001), which means that every 1 mmol/L increase of glucose level was associated with 12% lower odds of surviving. This finding is in agreement with previous studies that have shown that diabetes, which is associated with high glucose levels, is another COVID-19 risk-factor (*25*).

The role of serum albumin and its blood level in COVID-19 are not yet completely understood. Association of low albumin with poor outcomes is not unique to COVID-19: low levels are associated with inflammation (*30*), cancer (*31*), and are generally regarded as detrimental in many medical conditions (*32*). Hypoalbuminemia at triage has been found to predict increased 30-day mortality among all acutely admitted hospital patients (*33*) and to be a risk factor for Acute Respiratory Distress Syndrome (ARDS) (*28*). One potential cause for low albumin levels in these COVID-19 cases is prior comorbidities, such as malnutrition or liver disease, which predispose an affected patient to a more severe course of the illness. A second potential cause is that albumin levels decrease with age (**Fig. 3D**) (*34*). Lastly, it is possible that the disease itself can decrease albumin levels, e.g., by damaging the liver and impacting albumin production due to the virus attacking bile ducts and possibly other hepatic cells (*35*) or by hepatotoxicity of a COVID-19-induced cytokine storm (*36*). Our analysis shows only a minor albumin decline in patients that died (**Fig. 3C**), suggesting that this cause is unlikely to be a major one. However, it cannot be excluded that a significant drop in albumin levels happened in the early stages of COVID-19, prior to the patients’ admission to the hospital. This may have happened in Tongji Hospital, which admitted mostly severely ill patients. In any case, albumin levels may serve as a reliable predictor of COVID-19 severity. Serum albumin infusion aimed at correcting hypoalbuminemia, as used in severe liver damage, should be considered for COVID-19 patients (*37, 38*).

### Dexamethasone effectiveness on COVID-19 may be affected by hypoalbuminemia, drug-drug displacement, and diabetes

Our analysis of structural and clinical data demonstrates that serum albumin plasma level, competing drugs, and albumin glycation are important clinical variables that may influence the effectiveness of dexamethasone in treating COVID-19 patients. Each of these three factors decreases albumin’s binding capacity, which can have two adverse implications on dexamethasone blood transport: shorter half-life of the drug and a potential toxicity at its peak concentration. Average half-life of dexamethasone in pneumonia patients was reported to be 6.9 hours for 6 mg oral administration and 9.0 hours for 4 mg intravenous injection (*39*). Because albumin acts as a storage of drugs, the reduced binding capacity is likely to result in shorter half-life of dexamethasone, decreasing its therapeutic action. Moreover, reduced binding capacity increases the risk of dexamethasone toxicity because the maximum concentration of unbound dexamethasone increases. Side effects of dexamethasone are common, and low albumin levels has been observed to significantly increase the risk of its toxicity (*40, 41*). Therefore, clinicians should consider a modified dexamethasone regimen in COVID-19 treatment when: 1) patients suffer from hypoalbuminemia, 2) co-administered drugs compete with dexamethasone for DS7, and 3) DS7 is compromised by glycation due to diabetes. Notably, the presence of all three of these conditions may have a strong compounding effect. Dexamethasone regimen modifications could involve adjusting for patient’s weight and/or splitting the daily dose into multiples to achieve sufficient plasma concentration of dexamethasone over time while reducing possible peak toxicity, especially if higher than 6 mg doses are used (*42*). In other words, we call to flatten the dexamethasone concentration curve. Another feasible way of bypassing the limitations of vascular transport of dexamethasone would be to administer it via inhalers (*43*); it has been suggested that such steroid use may play a protective role in reducing vulnerability to COVID-19 (*44*). Once albumin levels of patients in the RECOVERY trial are released, a further analysis should be performed to establish whether there is a correlation between the effectiveness of dexamethasone intervention and hypoalbuminemia, drug co-administration, and serum albumin glycation.

## Conclusions

The presented molecular analysis of the binding of dexamethasone to serum albumin in combination with the risk-factors identified from clinical data offer promising strategies for maximizing dexamethasone effectiveness in the treatment of severe COVID-19. Adjusting the dexamethasone regimen may be necessary for patients with hypoalbuminemia, diabetes, or co-administration of large doses of drugs that bind to Drug Site 7. Moreover, oral inhalers should be evaluated as a better route of dexamethasone administration, which avoids the limitations of vascular transport. Our analysis also strengthens the case for the evaluation of albumin infusion aimed at correction of hypoalbuminemia. Guidelines for treatment of COVID-19 are still rapidly emerging and will remain critical even after a vaccine is developed, and dexamethasone will likely play a part in that treatment.

## Acknowledgments

We thank Alexander Wlodawer for his critical reading and discussions of the manuscript. We also thank Keith Brister, Zdzislaw Wawrzak, Spencer Anderson, and Joseph Brunzelle at LS-CAT sector 21 for their assistance in data collection.

## Author contributions

W.M, M.C., M.P.C, D.B., and I.G.S. conceived of the project and designed the research; K.A.M. purified the protein and determined the ESA-dexamethasone structure; M.P.C. and I.G.S. refined and analyzed the structure; D.B. conducted the statistical analyses of clinical data; I.G.S, M.P.C., and D.B., wrote the manuscript with active input from K.A.M., M.G., D.R.C., M.P., M.C., and W.M.

## Data and materials availability

Diffraction images were deposited to the Integrated Resource for Reproducibility in Macromolecular Crystallography at http://proteindiffraction.org with DOI: 10.18430/m3.irrmc.5571. Atomic coordinates and structure factors for the structure were deposited in the Protein Data Bank with accession code 6XK0. Reproducible statistical analysis and plotting scripts are available at https://github.com/dabrze/covid_albumin_levels.

## Supplementary Materials

Materials and Methods

Figures S1-S2

Tables S1-S4

References (*45–72*)

