## Supplementary Materials for "Molecular determinants of vascular transport of dexamethasone in COVID-19 therapy"

**This PDF file includes:**

Materials and Methods
Supplementary Text
Figs. S1 to S2
Tables S1 to S4

### Materials and Methods

#### Materials

ESA was purchased from Equitech-Bio (#ESA62; ≥96% purity; Kerrville, TX, USA) as lyophilized powder and purified further as described in the next paragraph. Dexamethasone was purchased from Sigma-Aldrich (#D1756; ≥98% purity; St. Louis, MO, USA). The purity of all reagents was reported by vendors.

### Protein purification and crystallization

ESA was dissolved in a buffer containing 10 mM Tris (pH 7.5) and 150 mM NaCl. Size exclusion chromatography using a Superdex 200 column attached to an ÄKTA FPLC (GE Healthcare) was used for further purification and to separate the dimeric and monomeric fractions of ESA. The purification buffer was the same as the buffer in which the protein was dissolved. The absorbance at 280 nm, measured with a Nanodrop 2000 (Thermo Scientific), was used to estimate protein concentrations using the extinction coefficient (ε280-ESA = 27,400 $M^{-1}$ $cm^{-1}$) and molecular weight ($MW_{ESA}$ = 65,700 Da). Collected fractions of monomeric ESA were concentrated to 15 mg/mL using an Amicon Ultra Centrifugal Filter (Millipore Sigma, #UFC903024) with a molecular weight cut-off (MWCO) of 30 kDa.

Protein crystallization was performed in 15-well hanging drop plates (EasyXtal 15-Well Tools, Qiagen). Prior to crystallization, dexamethasone powder in 10-fold molar excess was added to the concentrated protein solution (15 mg/mL ESA) in purification buffer. The mixture was incubated for 60 min at room temperature and then used for crystallization with some of the undissolved powder in suspension. Aliquots of 1 µL of the mixture were combined with 1 µL of reservoir solution (1.8M ammonium dihydrogen citrate at pH=7.0). Harvested crystals were flash-cooled in liquid nitrogen using a 1:1 mixture of Paratone® N and mineral oil as a cryoprotectant.

### Data collection and structure determination

Diffraction data were collected at 100 K at the 21 ID-F beamline of the Advanced Photon Source, Argonne National Laboratory. The collected data were processed, integrated, and scaled with HKL-3000 using corrections for radiation decay and anisotropic diffraction (*45*–*47*). The resolution cut-off and the number of images to be included in the final dataset were chosen based on the values of CC½, $\langle I \rangle / \langle \sigma(I) \rangle$, completeness, and $R_{meas}$. The initial phases were determined by molecular replacement with PDB ID: 3V08 as the template. The structure was refined with hydrogen atoms in riding positions using HKL-3000, seamlessly integrated with REFMAC (*45*, *46*) and other programs from the CCP4 package (*48*–*50*). Coot (*51*, *52*) was used for manual correction of the model. The protein model was placed in the standardized position in the unit cell using the ACHESYM server (*53*). TLS groups were determined and set up with a stand-alone version of the TLS Motion Determination server (*54*). The use of TLS parameters was justified by a significantly improved $R_{free}$ and the Hamilton R-factor ratio test (*55*) as implemented in HKL-3000. The structure refinement and model completion followed state-of-the-art, recently published guidelines (*56*, *57*), thus avoiding problems observed for some SARS-CoV-2 drug target models (*58*) . Both MOLPROBITY (*59*) and wwPDB validation servers (*60*) were used for model validation. PyMOL (The PyMOL Molecular Graphics System, Version 1.5.0.3, Schrödinger, LLC) was used for the preparation of structural figures. All experimental steps were tracked using LabDB (*61*). Molstack (*62*, *63*) was used for interactive visualization of the model and the electron density maps online. Diffraction images were deposited to the Integrated Resource for Reproducibility in Macromolecular Crystallography at http://proteindiffraction.org (*64*, *65*) with DOI: 10.18430/m3.irrmc.5571. Atomic coordinates and structure factors for the structure were deposited in the Protein Data Bank with accession code 6XK0. Statistics for diffraction data collection, structure refinement, and structure quality are presented in **Table S1.**

The raw diffraction dataset for the albumin-dexamethasone structure presented herein was collected nine years ago. The structure had to wait due to other pending projects, only to become more consequential now that it can be framed in the context of our recent work on albumin ligands and the albumin level data from Wuhan COVID-19 patients. This example shows that sometimes the importance of basic science experiments is not immediately apparent, but the questions they answer can one day become of tremendous value.The use of LabDB (*61*) was essential for locating this dataset, demonstrating that a reliable database with extensvive descriptions of experiments has a vital advantage for research labs.

**Clinical data of COVID-19 patients**

Albumin and glucose levels of patients from Tongji Hospital, Wuhan, China were taken from the dataset published by Li Yan *et al.* (*26*), available at https://github.com/HAIRLAB/Pre_Surv_COVID_19. The dataset was made public under the MIT License. As described in (*26*), the blood test results of the patients were collected between January 10 and February 18, 2020. The original dataset describes 375 patients, but here we analyze only the 373 patients for whom albumin levels were reported. Most of these patients (356 out of 373) had multiple blood samples taken throughout their stay in the hospital. If not stated otherwise in the text, we used the last sample taken to calculate statistics, as it most accurately matches the patient's outcome (death or survived). However, the median changes of patients' albumin levels over time were very small (**Fig. 3C**), and statistics obtained using other samples would be almost identical. Reproducible analysis scripts are available at https://github.com/dabrze/covid_albumin_levels.

**Statistical methods**

The differences in sample means were assessed using two-tailed Welch's t-tests at significance level $\alpha = 0.05$. Prior to assessing the significance of the differences, the samples were checked for normality using the Kolmogorov-Smirnov test with $\alpha = 0.05$ and visually inspected using Q-Q plots. For calculating the correlation between admission and final albumin levels, we used the Pearson product-moment correlation coefficient, two-tail-tested against a $t$ distribution with $n$-2 degrees of freedom ($df_{Died} = 346$, $df_{Survived} = 396$) at $\alpha = 0.05$; patient's with only one albumin level available were excluded from the calculation.

Logistic regression models were used to estimate the association between patient albumin levels and their survival or death. The model was adjusted to take into account such confounding factors as age, gender, glucose levels. All confounders were checked for potential effect modification, but no effect modification was found as all interaction terms exhibited $p > 0.2$. Detailed odds ratios with confidence intervals and $p$-values are presented in **Table S3**.

### Supplementary Text

Overall structure of ESA, dexamethasone conformation, and other molecules bound

During model building, the polypeptide chain was almost entirely completed, except for the first three N-terminal residues, which were not located in the electron density maps. Albumin consists of three homologous domains: I (residues 1–195), II (196–383), and III (384–585); each domain contains two subdomains (A and B) composed of 6 and 4 alpha-helices, respectively. The overall fold of ESA in this structure is essentially identical to previously published structures of ESA and HSA (**Table S2**). According to the Dali server (*66*), the closest structure is the ESA ligand-free structure (PDB ID: 3V08) with RMSD 0.4 Å.

During the refinement, a different orientation of dexamethasone was tried as an alternative, in which the drug was rotated by 180° along the axis perpendicular to its rings. However, in the alternative orientation one of the four rings of dexamethasone was not covered by strong electron density, and the compound did not form any hydrogen bonds with the protein, clearly supporting the chosen conformation.

In addition to the dexamethasone, one citrate molecule and one fatty acid molecule were located in the structure. The citrate molecule is located inside the cleft between domains I and III, near DS9, at the same position as in ESA-testosterone complex; citrate was a major component of the crystallization cocktail. The fatty acid molecule, which was likely retained during purification of ESA from blood, is located in fatty acid site 8 (FA8). Very week electron density is also observed in DS4, which does not allow for any certainty in the interpretation. This density was accounted for by four UNX atoms.

<u>Conservation of Drug Site 7 between HSA and ESA</u>

Fourteen out of 15 residues involved in dexamethasone binding to ESA are conserved in HSA (**Figure S1**). Only one residue is different: Ala481 in ESA corresponds to Val in HSA. This small hydrophobic-for-hydrophobic difference is unlikely to affect dexamethasone binding in HSA because it is a kind-for-kind change that does not introduce any clashes with the ligand. In fact, in the two structures of HSA that have a ligand bound to this site in the vicinity of this residue – complexes with diclofencac (PDB ID: 4Z69) and ibuprofen (PDB ID: 2BXG) – the Val residue turns away from the ligand. This Val conformation makes its CB atom to be the closest atom to the ligand, thus making Val very similar to Ala in terms of distances to the ligand. Therefore, the conservation of amino acid residues in DS7, which leads to the essentially identical hydrophobic environments, suggests that dexamethasone binds to HSA in the same site as in ESA.

<u>Glycation of residues forming Drug Site 7</u>

HSA has multiple Lys and Arg residues that are known to be glycated (*67–70*). Depending on the method, it is estimated that up to 6% of the HSA in a healthy human is glycated, while in diabetic patients these values are 2-5 times higher (*67*, *69*, *71*). Residues of DS7 residues that are likely to undergo glycation are Arg208, Lys211, Lys350 (Arg209, Lys212, and Lys351, respectively, in HSA) (*67*). Glycation of Lys211 or Lys350 will not necessarily cause a disruption of dexamethasone binding because these residues are pointed outside of the binding site, towards the solution (Fig. S2B). However, glycation of Arg208 is likely to prevent drug binding in this site due to potential steric clashes and disruption of the binding site due to possible elimination of the Arg208-Asp323 salt bridge, thus decreasing SA binding capacity.

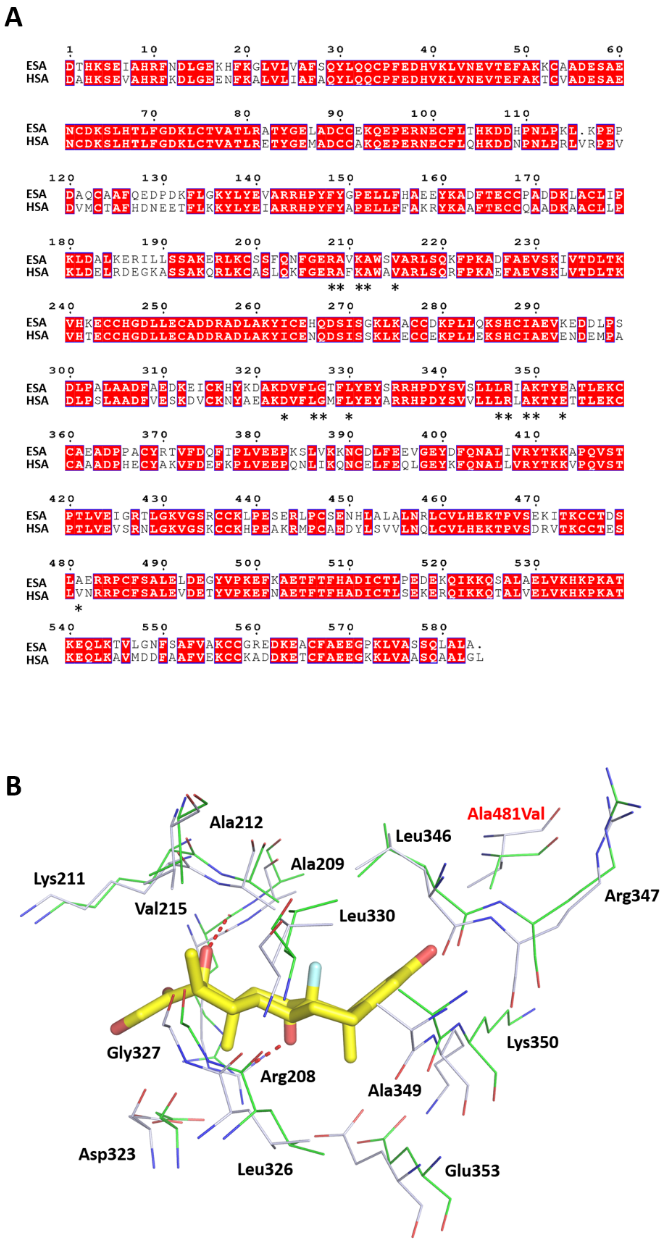


**Fig. S1.**

**Comparison of ESA and HSA.** (**A**) Alignment of ESA and HSA sequences. Identical residues are highlighted in red; residues involved in dexamethasone binding to DS7 in ESA and analogous residues in HSA are marked with stars. (**B**) Superposition of DS7 binding dexamethasone in ESA (PDB ID: 6XK0) and analogous site in HSA (PDB ID: 4K2C). Dexamethasone is shown in stick representation with carbon atoms in yellow, oxygen atoms in red and fluoride atom in cyan. Residues within 5 Å are shown in line representation, residues from ESA are shown with carbon atoms in green while residues from HSA are shown with carbon atoms in gray, oxygen and nitrogen atoms are red and blue, respectively. Residue numbers correspond to positions in ESA. Ala481 is the only residue at this site that differ in HSA (Val in HSA). Hydrogen bonds are shown as red dashes.

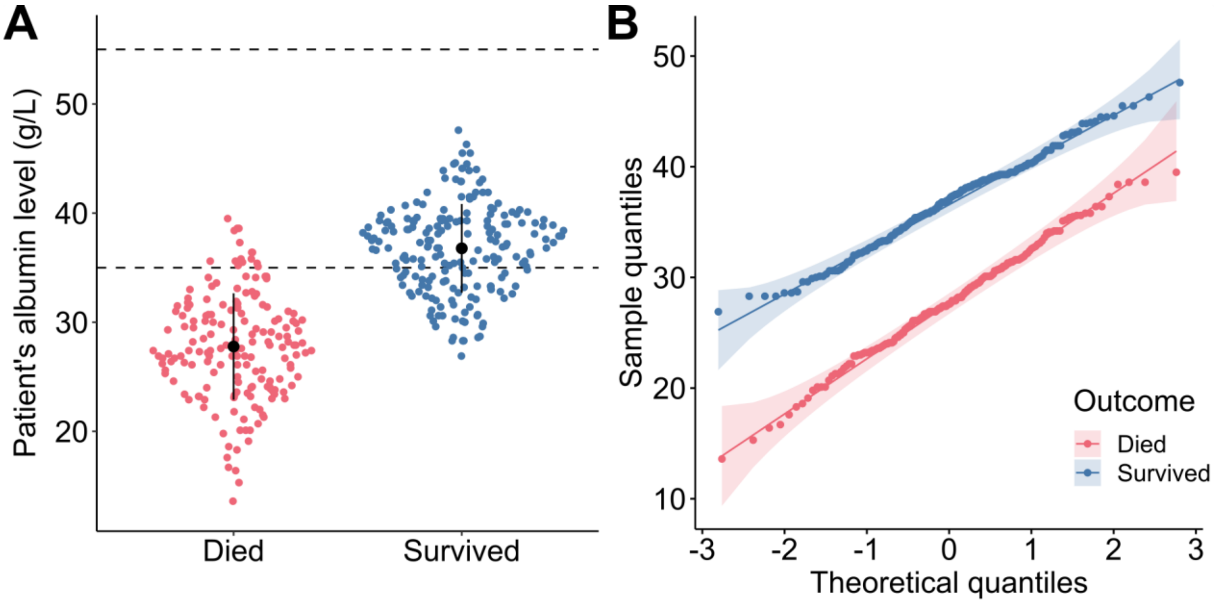


**Fig. S2.**

**Distribution of COVID-19 patients grouped by outcome.** (**A**) Violin strip charts (sina plots) of patients that died (red) and (survived), after being admitted to Tongji Hospital. (**B**) Q-Q plot for the albumin levels of patients that died and survived COVID-19. The shaded areas represent 95% confidence intervals.

**Table S1.**

Data collection, structure refinement, and structure quality statistics.

| | |
|---|---|
| **PDB ID:** 6XK0 | |
| **Diffraction data DOI:** 10.18430/m3.irrmc.5571 | |
| **Data collection statistics** | |
| **Resolution (Å)** | 50.00-2.40 (2.44-2.40) |
| **Beamline** | 21-ID-F |
| **Wavelength (Å)** | 0.979 |
| **Space group** | $P6_1$ |
| **Unit-cell dimensions (Å)** | a=b=95.0 c=143.6 |
| **Protein chains in the ASU** | 1 |
| **Completeness (%)** | 99.8 (98.3) |
| **Number of unique reflections** | 28870 (1412) |
| **Redundancy** | 4.9 (3.5) |
| **$\langle I \rangle / \langle \sigma(I) \rangle$** | 15.3 (1.3) |
| **CC ½** | (0.61) |
| **$R_{merge}$** | 0.110 (1.053) |
| **$R_{meas}$** | 0.123 (1.221) |
| **Refinement statistics** | |
| **$R_{work}/R_{free}$** | 0.203/0.249 |
| **Bond lengths rmsd (Å)** | 0.002 |
| **Bond angles rmsd (°)** | 1.1 |
| **Mean ADP (Å$^2$)** | 57 |
| **Mean ADP for dexamethasone (Å$^2$)** | 90.5 |
| **Number of protein atoms** | 4589 |
| **Mean ADP for protein (Å$^2$)** | 58 |
| **Number of water molecules** | 228 |
| **Mean ADP for water molecules (Å$^2$)** | 39 |
| **Clashscore** | 2.71 |
| **MolProbity score** | 1.25 |
| **Rotamer outliers (%)** | 0.60 |
| **Ramachandran outliers (%)** | 0.00 |
| **Ramachandran favored (%)** | 96.88 |

Values in parentheses are for the highest resolution shell. Ramachandran plot statistics are calculated by MolProbity.

**Table S2.**

RMSD values [Å] for aligned Cα atoms of ESA-steroid complexes and ligand-free ESA and HSA structures.

| | ESA-dexamethasone (PDB ID: 6XK0) | ESA-testosterone (PDB ID: 6MDQ) | ESA-ligand free (PDB ID: 3V08) | HSA-ligand free (PDB ID: 4K2C) |
|---|---|---|---|---|
| ESA-dexamethasone (PDB ID: 6XK0 ) | - | 1.1 | 0.4 | 1.6 |
| ESA-testosterone (PDB ID: 6MDQ) | 1.1 | - | 1.1 | 1.6 |
| ESA-ligand free (PDB ID: 3V08) | 0.4 | 1.1 | - | 1.7 |
| HSA-ligand free (PDB ID: 4K2C) | 1.6 | 1.6 | 1.7 | - |

**Table S3.**

Associations between gender, age, glucose level, albumin level and survival or death in univariate and multivariate logistic regression analyses.

| **Analysis** | **Odds ratio of survival (95% CI)** | **p-values** |
|---|---|---|
| Univariate analysis (unadjusted models) | | |
| Albumin level | 1.56 (1.44–1.71) | $p < 0.001$ |
| Female | 2.82 (1.83–4.37) | $p < 0.001$ |
| Age | 0.90 (0.88–0.92) | $p < 0.001$ |
| Glucose level | 0.76 (0.70–0.81) | $p < 0.001$ |
| Multivariate analysis (adjusted model) | | |
| Albumin level | 1.51 (1.37–1.69) | $p < 0.001$ |
| Female | 2.19 (1.04–4.72) | $p < 0.05$ |
| Age | 0.92 (0.89–0.94) | $p < 0.001$ |
| Glucose level | 0.89 (0.81–0.96) | $p < 0.01$ |

**Table S4.**

Comparison of mean (s.d) of albumin levels and patient outcomes in different studies. N denotes the sample size of the study. ICU: Intensive Care Unit, ARDS: Acute Respiratory Distress Syndrome.

| Country | N | Group 1 (g/L) | Group 2 (g/L) | Study |
|---|---|---|---|---|
| Spain | 48 | ICU: 29.0 (5.2) | Non-ICU: 39.2 (4.2) | (*29*) |
| China[*] | 21 | Severe: 29.6 (28.6–33.0) | Moderate: 37.2 (35.8–38.8) | (*27*) |
| China[*] | 201 | ARDS: 30.4 (27.15–33.35) | Non-ARDS: 33.7 (30.95–36.30) | (*28*) |
| China[*†] | 2623 | Died: 31.1 (27.9–34.2) | Non-/critical: 36.6 (33.2–40.4)/32.2 (29.6–35.7) | (*36*) |
| China[‡] | 910 | Severe: 35.0 (2.4) | Non-severe: 40.5 (2.2) | (*72*) |
| China | 373 | Died: 27.8 (4.9) | Survived: 36.8 (4.1) | current |

[*]Albumin level characterized by the median and quartiles (Q1–Q3); [†]Albumin levels measured at admission; [‡]Meta-analysis of data form different hospitals
